# Patient-Derived Glioma Models Preserve Tumor Heterogeneity and Identify Stearoyl-CoA Desaturase1 (SCD1) as a Candidate Biomarker for Precision Immunotherapy

**DOI:** 10.64898/2026.09.01.748569

**Authors:** Margarita Gutova, Eric Ma, Heini N. Natri, Sean Sepulveda, Sharvari Mankame, Spencer Uyematsu, Jonathan Hibbard, Maryam Aftabizadeh, Xiujian Ma, Renate Starr, Brenda Aguilar, Julie Ostberg, Lisa Feldman, Massimo D’Apuzzo, Nicholas E. Banovich, Floris Barthel, Behnam Badie, Christine E. Brown

## Abstract

**Background:** Pediatric and adult brain tumors, including glioblastoma, astrocytoma, ependymoma, and medulloblastoma, remain associated with poor prognosis despite advances in surgery, radiation, and chemotherapy. Therapeutic resistance, tumor heterogeneity, and treatment-related toxicity highlight the need for clinically relevant models that enable precision medicine and immunotherapy development.

**Methods:** Freshly dispersed tumors (FDTs), low-passage patient-derived brain tumor (PBT) spheroid lines, and matched patient-derived xenograft (PDX) models were established from patients with primary brain tumors. Models were characterized using single-cell and bulk RNA sequencing, whole-exome sequencing, multiparameter flow cytometry, and immunohistochemistry. PBTs were compared with matched FDTs to evaluate model fidelity.

**Results:** PBT lines were established in approximately 68% of cases and retained key patient-specific genomic alterations, including IDH1, MGMT, TP53, and PTEN, with expression of therapeutically relevant targets, including IL13Rα2, EGFR, HER2, WNT1, JAK1/2, and NOTCH1-4. Gene expression profiles of PBTs closely correlated with matched FDTs (R = 0.38, P = 0.0038). PBT and PDX models preserved intratumoral heterogeneity and non-clonal populations, enabling identification of therapy-resistant subclones during *in vitro* selection. Molecular analyses identified Stearoyl-CoA Desaturase1 (SCD1) as an overexpressed biomarker across glioma PBTs and matched patient tumors. Ingenuity Pathway Analysis identified SCD1 as an upstream regulator of EGFR-, TP53-, and CYCS-associated signaling networks implicated in tumor progression and immune suppression.

**Conclusions:** Clinically relevant PBT and matched PDX models recapitulate molecular, transcriptional, and histopathological characteristics of primary brain tumors. These platforms provide tools for biomarker discovery, therapeutic testing, and precision immunotherapy development, while identifying SCD1 as a biomarker and therapeutic target in glioma.

## Introduction

High-grade gliomas (HGGs), including glioblastoma (GBM), are among the most challenging and incurable brain tumors^1–4^. The refractory nature of GBM, coupled with side effects of current treatments, such as surgery, radiation, chemotherapy, and electric fields, underscores the pressing need for more effective therapeutic strategies^5–9^. A significant hurdle in developing more effective therapies, including immunotherapies, is the lack of preclinical models that accurately capture the complex cellular and genetic heterogeneity observed between and within patient tumors^10, 11^. Although current standard treatments provide some benefit, they often fall short due to tumor heterogeneity and the varied responses of individual patients ^7, 12, 13^. Over the past two decades, GBM research has established patient-derived glioma stem cell (GSC) and spheroid cultures as biologically relevant models that preserve key features of the parental tumors^14^. Early studies demonstrated the existence of self-renewing, tumor-initiating cancer stem cells and showed that serum-free culture conditions better maintain tumor genotype, gene expression, and phenotypic characteristics than conventional serum-grown cell lines^15, 16^. Subsequent work expanded these approaches by developing scalable culture systems, large patient-derived biobanks, and molecularly diverse reference panels, while confirming the preservation of histologic, genomic, transcriptional, and epigenetic features ^17^. More recent studies have further linked transcriptional states to therapeutic vulnerabilities and highlighted culture-associated evolutionary changes^18^. Despite these advances, relatively few studies have comprehensively compared freshly dissociated tumors with their matched low-passage cultures using integrated genomic, transcriptional, and cellular analyses, leaving important questions regarding model fidelity and translational applicability.

We hypothesize that personalized therapy, tailored to the unique molecular and cellular profiles of each patient’s tumor, will improve treatment outcomes^19^. To address this, we have developed patient-derived brain tumor (PBT) low-passage cell lines derived from freshly dispersed tumors (FDTs) and developed patient-derived xenograft (PDX) models that closely mimic human glioma development and biomarker expression. These models provide a more accurate representation of human GBM and HGGs, allowing for a comprehensive evaluation of immunotherapies and other targeted treatments.

Our approach involves generating PBT spheroid lines *in vitro* and PDX models *in vivo* from multiple patient tumors, enabling the empirical identification of human tumor-specific characteristics and therapeutic targets that may not be apparent from genomic or transcriptomic analyses alone^20^. The variability in mutations and tumor characteristics necessitates a more detailed understanding of intra-tumoral heterogeneity and patient-to-patient differences, which can impact therapeutic response. We have addressed this need by creating a patient-specific cohort of PBT spheroids with detailed clinical, molecular, and phenotypic data to evaluate differential responses and resistance to CAR T cell therapy. Our platform integrates clinical data (including sex, age, diagnosis, and patient history) with tumor pathological features, allowing for a more precise assessment of therapeutic efficacy. This innovative platform, which combines patient-specific tumor models, offers a unique tool for evaluating individual tumor responses to CAR T cells and other therapies^13, 21–23^. This platform also facilitates the exploration of heterogeneity in treatment responses across multiple patient tumors^24–26^. By identifying the underlying factors contributing to response variability, we aim to guide the development of next-generation therapeutic strategies and combinational treatments.

## Methods

### Tumor Sample Collection and Analysis Protocol

#### Sample Collection

Tumor samples were collected from glioma patients who participated in clinical trials at COH (IRB#13384) immediately following their surgical procedures, ensuring that the samples were as fresh and representative of the patient’s tumor as possible^27, 28^.

#### Molecular Profiling

The HopeSeq Glioma Comprehensive Panel was used to detect a range of genetic alterations, including mutations, gene fusions, and copy number variation (CNV) commonly associated with gliomas. The panel was also used to assess critical biomarkers for treatment response such as p53, IDH1/2, TERT, ATRX, CDKN2A, EGFRvIII, and O6-methylguanine-DNA methyltransferase [MGMT] gene methylation status^29^.

#### Single Cell RNA-Sequencing (scRNA-seq)

scRNA-seq was performed on FDT samples to capture and analyze gene expression at the individual cell level. The scRNA-seq data leveraged here comprises 41 cryopreserved HGG FDT samples, including 29 GBM samples. ScRNA-seq was performed using the 10× Chromium platform. Briefly, samples were multiplexed and sequenced using either whole exome genotype-based demultiplexing, antibody-based demultiplexing, or sequenced without pooling. 60,000 cells were loaded onto each Gel Bead-in-Emulsion (GEM) reaction. Library preparation was performed according to manufacturer protocols and sequenced on Illumina NovaSeq6000 at the recommended depth. Sequence data were processed using 10× Genomics Cell Ranger V5.0 and Ensemble 98 and analyzed using Seurat v5^30^. Cell types were annotated based on canonical marker gene expression. Using the scGSVA R package (https://github.com/guokai8/scGSVA), we analyzed data from each tumor for the enrichment of gene expression signatures associated with GBM meta-modules ^31^. For comparison with PBT cell lines, pseudo-bulk profiles of the malignant cells in the FDT samples were generated by calculating the average log_2_(transcripts per million [TPM]) for each gene.

#### Clinical Data Collection

Comprehensive, de-identified clinical information was collected for each patient and is summarized in Table 1. This dataset includes key demographic and clinical variables such as age, sex, diagnosis, treatment history, and clinical outcomes. These contextual data provide critical insights into the biological and clinical relevance of the tumor samples and enable meaningful correlations between molecular alterations and patient-specific therapeutic responses and disease progression.

**Table 1.**
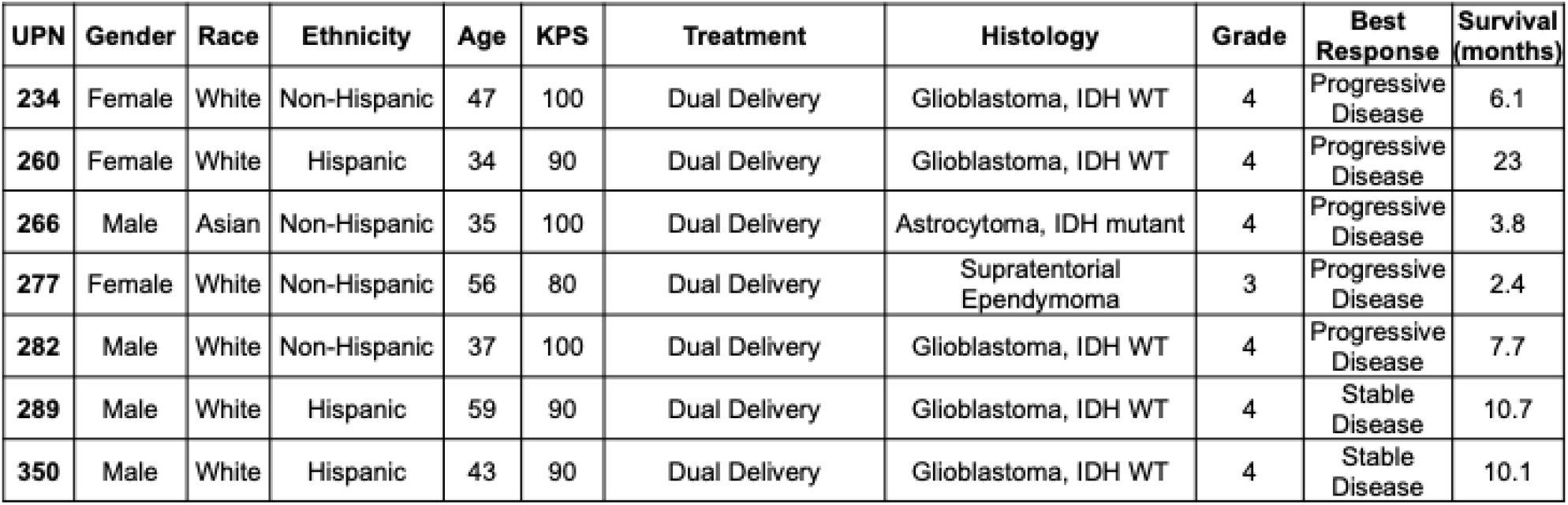
Patient Demographics.

#### Cell Line Establishment

Tumor samples were transferred to the COH CAR T Cell group for the establishment of PBT cell lines. Tumor neurosphere formation was performed as previously described ^19, 32, 33^. Briefly, glioma neurosphere culture protocol was adapted from Brown et al., 2009^33^. Patient-derived glioma specimens, graded per WHO guidelines, were obtained under IRB-approved protocols and processed for *in vivo* and *in vitro* modeling. Tumor tissues were minced and either implanted subcutaneously into NOD-scid mice or enzymatically dissociated into single cells using Tumor Dissociation Kit, human (Miltenyi, 130-095-929). Tumor spheres (TSs) were cultured in neural stem cell medium [DMEM: F12 supplemented with B27, heparin, L-glutamine, epidermal growth factor (EGF [20 ng/mL]) and basic fibroblast growth factor (bFGF [20 ng/mL]).

#### PBT Cell Culture Propagation Protocol

Each PBT cell line was seeded at densities ranging from 25,000 to 55,000 cells/cm² in T75 flasks. Cell counts were performed at each passage, which occurred every 3 to 7 days, using the Muse™ Personal Cell Analysis System (Catalog No: **0500-3115**). The growth rate of each cell line was calculated using the formula provided in Equation 1 ^34^. Additionally, the time and date of each passage were meticulously recorded to determine the doubling time of the cells, calculated as shown in Equation 2. This approach ensured precise monitoring of cell proliferation and provided valuable data on the growth kinetics of each PBT line.

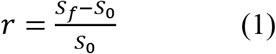

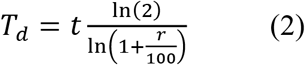

*r* = Growth Rate
*S*_*f*_ = Final Count
*S*_0_ = Seeding Density
*T*_*d*_ = Doubling Time
*t* = Time

#### Preparation of Single-Cell Suspensions from Adherent and Suspension Neural Cell Lines

*<u>For adherent cell lines</u>*, culture media were aspirated, and cells were rinsed with 5 mL of phosphate-buffered saline (PBS). Cells were then detached using 3 mL of Accutase (Stem Cell Technologies, Catalog No. NC1793126) and incubated for 5 min at room temperature. Following detachment, 9 mL of complete media was added to neutralize the enzymatic activity of Accutase. The resulting cell suspension was filtered through a 40 µm strainer to obtain a single-cell suspension, and a 25 µL aliquot was taken for cell counting.

For *<u>suspension cell</u> lines* such as neurospheres, culture media containing neurospheres were transferred into a 50 mL conical tube and centrifuged to pellet the cells. After discarding the supernatant, the neurosphere pellet was resuspended in 1 mL of Accutase and incubated to facilitate dissociation into single cells. Accutase activity was quenched by adding 3 mL of complete media, and the suspension was passed through a 40 µm filter. A 25 µL aliquot of the filtered suspension was used for cell counting. Cell line images were captured using a Zeiss Axio Observer Z1 Microscope at 10× magnification. The images were obtained from 96-well plates 24 h post-passaging, adhering to the previously described cell density.

#### Flow Cytometry Analysis of PBT Lines

Cells were cultured without EGF and FGF one passage prior to flow cytometry analysis to prevent interference with EGF receptor (EGFR) staining. Single-cell suspensions of PBT lines were prepared by washing with 100 µL of HBSS supplemented with 2% fetal calf serum (FCS) and sodium azide (FSS), then centrifuged at 800 × g for 2.5 min. The supernatants were discarded, and the cells were incubated with one of three distinct antibody panels: PBT flow cytometry panel, Stem panel, and Miscellaneous panel for 30 min at 4 °C. After incubation, the cells were washed and spun down twice with FSS and resuspended in 100 µL of FSS. Just before analysis on the MASCQuant Analyzer 10 (Miltenyi), 50 µL of DAPI were added to the cell suspension to enable viability staining.

The <u>PBT flow cytometry panel</u> included markers such as CD73 (BD Biosciences, 563198), Her2 (Biolegend, 324404), IL13Rα2 (Biolegend, 354404), B7H3 (Biolegend, 351010), EGFR (Biolegend, 352910), and CD39 (Biolegend, 328226). Chlorotoxin (CLTX), used in the panel, was synthesized at COH, with Cy5.5 NHS Ester (GE Healthcare, PA15601) and Slide-A-Lyzer MINI (Thermo Fisher Scientific, 69558) used for labeling.

The <u>Stem panel</u> contained CD90 (Biolegend, 328126), CD133 (Miltenyi, 130-113-111 and 130-113-188), CD15 (Biolegend, 323006), CD56 (Catalog number 560842, BD Biosciences), and CD44 (BD Biosciences, 560532), targeting stem cell-related markers.

The <u>Miscellaneous panel</u> contained a range of additional markers, including PDL1 (Biolegend, 329734), CD70 (BD Biosciences, 555834), CD119 (Invitrogen, 12-1199-42), HLABC (BD Biosciences, 555554), HLADRDPDQ (Biolegend, 361708), CD304 (Biolegend, 354505), and CD54 (Biolegend, 353122).

#### OncoScan® Assay Protocol

##### Sample Preparation

DNA was extracted from each PBT line using the Maxwell® 16 FFPE Plus LEV DNA Purification Kit (Promega), adhering strictly to the manufacturer’s instructions to ensure high-quality DNA extraction.

##### DNA Quantification

DNA concentration was measured using the Quantus® Fluorometer (Promega) to ensure accurate quantification for the success of subsequent hybridization and analysis steps.

##### Hybridization and Processing

To detect copy number variation across the genome, DNA samples were prepared and hybridized on the OncoScan® assay array following the detailed protocol provided by Affymetrix.

##### Array Features

The OncoScan® assay array includes over 220,000 single nucleotide polymorphisms, providing a genome-wide resolution of 300 kb and a higher resolution of 50– 100 kb in approximately 900 cancer-related genes. This high-resolution capability was used for detailed genomic analysis of PBT lines.

##### Data Analysis

Nexus software was used to analyze the OncoScan® assay data, facilitating the identification and interpretation of CNV across the genome. We specifically focused on identifying chromosomal alterations, including gains or losses in key genomic regions in PBT lines. This analysis provided insight into the genomic mutations of the PBT lines, similarities with FDT samples, and the potential drivers of tumorigenesis.

#### Cell Line Selection

Cell lines were selected for further analysis based on their ability to form neurospheres and the availability of scRNA-seq data. Consequently, lines that were not part of the initial patient group or lacked RNA sequencing data were not included in the subsequent analysis group to ensure comprehensive evaluation of selected FDTs and PBTs.

Bulk **RNA sequencing** was performed on seven low-passage PBT cell line samples. Total RNA was extracted using the **RNeasy Mini Kit (Qiagen, 74104)**, and RNA integrity was confirmed with an Agilent Bioanalyzer (**RIN ≥ 8.0**). RNA-seq libraries were prepared using the **KAPA mRNA HyperPrep Kit** with poly(A) selection to enrich for mRNA and preserve strand specificity. Libraries were pooled and sequenced on an Illumina **NovaSeq 6000** platform, generating paired-end 2 × 108 bp reads, yielding approximately 20–36 million read pairs per sample. We used the R package signscore to analyze bulk-RNAseq profiles for GBM meta-module signatures^31^. A composite score (totalScore) was calculated by summing the enrichment scores for both up-regulated and down-regulated genes within each gene set^35, 36^.

#### HopeSeq Glioma Comprehensive Assay

The HopeSeq Glioma assay identifies prevalent glioma mutations associated with approved treatments, experimental therapies in clinical trials, or other relevant clinical implications. For this study, the HopeSeq Glioma assay was applied to FDT samples from patients immediately following their surgical procedures.

#### Hotspot Mutation Analysis of 87 Genes related to identification of targetable mutations

AKT1, AKT2, AKT3, ALK, AR, ARAF, AXL, BRAF, BTK, CBL, CCND1, CDK4, CDK6, CHEK2, CSF1R, CTNNB1, DDR2, EGFR, ERBB2, ERBB3, ERBB4, ERCC2, ESR1, EZH2, FGFR1, FGFR2, FGFR3, FGFR4, FLT3, FOXL2, GATA2, GNA11, GNAQ, GNAS, H3F3A, HIST1H3B, HNF1A, HRAS, IDH1, IDH2, JAK1, JAK2, JAK3, KDR, KIT, KNSTRN, KRAS, MAGOH, MAP2K1, MAP2K2, MAP2K4, MAPK1, MAX, MDM4, MED12, MET, MTOR, MYC, MYCN, MYD88, NFE2L2, NRAS, NTRK1, NTRK2, NTRK3, PDGFRA, PDGFRB, PIK3CA, PIK3CB, PPP2R1A, PTPN11, RAC1, RAF1, RET, RHEB, RHOA, ROS1, SF3B1, SMAD4, SMO, SPOP, SRC, STAT3, TERT, TOP1, U2AF1, XPO1

#### Full-length Mutation Analysis of 48 Genes

ARID1A, ATM, ATR, ATRX, BAP1, BRCA1, BRCA2, CDK12, CDKN1B, CDKN2A, CDKN2B, CHEK1, CREBBP, FANCA, FANCD2, FANCI, FBXW7, MLH1, MRE11A, MSH2, MSH6, NBN, NF1, NF2, NOTCH1, NOTCH2, NOTCH3, PALB2, PIK3R1, PMS2, POLE, PTCH1, PTEN, RAD50, RAD51, RAD51B, RAD51C, RAD51D, RB1, RNF43, SETD2, SLX4, SMARCA4, SMARCB1, STK11, TP53, TSC1, TSC2

#### Fusion Analysis of 51 Genes

AKT2, ALK, AR, AXL, BRAF, BRCA1, BRCA2, CDKN2A, EGFR, ERBB2, ERBB4, ERG, ESR1, ETV1, ETV4, ETV5, FGFR1, FGFR2, FGFR3, FGR, FLT3, JAK2, KRAS, MDM4, MET, MYB, MYBL1, NF1, NOTCH1, NOTCH4, NRG1, NTRK1, NTRK2, NTRK3, NUTM1, PDGFRA, PDGFRB, PIK3CA, PPARG, PRKACA, PRKACB, PTEN, RAD51B, RAF1, RB1, RELA, RET, ROS1, RSPO2, RSPO3, TERT

#### Copy Number Analysis of 47 Genes

AKT1, AKT2, AKT3, ALK, AR, AXL, BRAF, CCND1, CCND2, CCND3, CCNE1, CDK2, CDK4, CDK6, CDKN2A, CDKN2B, EGFR, ERBB2, ESR1, FGF3, FGF19, FGFR1, FGFR2, FGFR3, FGFR4, FLT3, IGF1R, KIT, KRAS, MDM2, MDM4, MET, MYC, MYCL, MYCN, NTRK1, NTRK2, NTRK3, PDGFRA, PDGFRB, PIK3CA, PIK3CB, PPARG, RICTOR, TERT, TSC1, TSC2

#### Generation of PDX Models of Glioma

PDX models were generated by intracranial engraftment of low-passage PBT lines into NOD.Cg-Prkdc^scid Il2rg^tm1Wjl (NSG) mice. NSG mice provide a severely immunodeficient background that supports efficient engraftment of human brain tumors and the development of clinically relevant PDX models. Following implantation, mice were monitored to assess tumor take and latency. Mice were anesthetized using a combination of ketamine/xylazine and maintained under gaseous isoflurane. A suspension of 0.1 × 10⁶ PBT cells in 2 µL of medium was stereotactically implanted into right frontal lobe, as described previously^37^. This technique ensured accurate and reproducible delivery of tumor cells into the brain parenchyma. Mice were monitored daily for clinical signs of tumor progression, including changes in body weight, grooming behavior, posture, and motor coordination. Upon the onset of neurological symptoms or significant weight loss indicative of tumor burden, animals were humanely euthanized. Brains were then collected for comprehensive histopathological analysis to confirm tumor establishment and assess histological features.

#### DNA Extraction and Library Preparation

Frozen tumor tissue and matched peripheral blood mononuclear cells (PBMCs, normal) were obtained from each patient. Genomic DNA (gDNA) was isolated using the NEB Monarch kits for Tissue (NEB #T3060) and for Cells (NEB #T3050L), which employ glass bead-based disruption to recover high molecular weight DNA. DNA concentration and integrity were assessed using the Qubit 1× dsDNA Broad Range Assay (Thermo Fisher Scientific, Waltham, MA, USA) and the Agilent TapeStation gDNA ScreenTape system (Agilent Technologies, Santa Clara, CA, USA). Indexed sequencing libraries were generated with the Twist EF 2.0 Library Preparation Kit (Twist Bioscience, South San Francisco, CA, USA), incorporating enzymatic fragmentation, end-repair, A-tailing, adapter ligation, and PCR amplification steps. A subset of each library was allocated for whole genome sequencing (WGS), while the remaining portion was used for targeted enrichment to allow complementary genome-wide and gene-focused analyses.

#### Targeted Sequencing Next Generation Sequencing

Target enrichment was performed using the Twist Fast Hybridization Target Enrichment protocol (Twist Bioscience). Up to eight indexed libraries were pooled for enrichment with a dual-panel design, including a mitochondrial panel and a custom glioma-specific panel targeting recurrently mutated genes in GBM. Libraries were normalized to equal concentrations and combined to yield a total of 1.5 µg DNA. The pooled mixture was vacuum-dried and hybridized with the capture probes at 60 °C for 4 h. Captured DNA fragments were retrieved using streptavidin-coated magnetic beads, followed by a series of washes and elution in 45 µL of molecular-grade water. Final enriched libraries were quantified using Qubit and TapeStation prior to sequencing. All libraries (tumor gDNA, normal genomic DNA) were sequenced on the Illumina NovaSeq X platform with 2 × 150 bp paired-end reads.

#### Variant Calling Analysis

Sequencing data from all cfDNA and gDNA libraries were processed using a custom pipeline incorporating fgbio and GATK tools optimized for UMI-based error correction. Adapter and empty reads were removed from raw FASTQ files using Cutadapt (v4.8), and the cleaned reads were aligned to the hg38 reference genome using BWA (v0.7.18). PCR duplicates were identified and collapsed into single consensus reads with fgbio (v2.2.1) to minimize sequencing artifacts. Base quality recalibration and somatic variant calling were performed using GATK Mutect2, followed by filtration of low-confidence and germline variants. High-confidence variants were functionally annotated using GATK Funcotator. The mean sequencing depth was approximately 800× for tumor samples and 1,200× for matched normal controls.

#### Immunohistochemical (IHC) Staining

was performed at the COH Pathology Core to evaluate the expression of specific tumor markers. The antibodies used included B7H3 (Rabbit monoclonal antibody, #Cat:14058, Cell Signaling Technology, diluted at 1:40), CD70 (Cat number:69209, Cell Signaling Technology, diluted at 1:50), Her2 (Rabbit monoclonal antibody, #Cat:2165, Cell Signaling Technology, diluted at 1:100), IL13Rα2 (Rabbit monoclonal antibody, #Cat:85677, Cell Signaling Technology, diluted at 1:100), PDL2 (Rabbit monoclonal antibody, #Cat:82723, Cell Signaling Technology, diluted at 1:25), pSMAD2 (Rabbit monoclonal antibody, #Cat:44-244G, Invitrogen, diluted at 1:100), and GD2 (Mouse monoclonal antibody, #Cat:554272, BD Pharmingen, diluted at 1:50). These antibodies were chosen to accurately profile the expression of biomarkers in tumor samples, providing valuable insight into their distribution and levels, which is crucial for understanding the biological characteristics of the tumors and evaluating potential therapeutic targets. IHC slides were scanned at 10× magnification using the NanoZoomer 2.0-HT slide scanner and analyzed with QuPath software, allowing for the quantification of total cell count, positive cell detection, positive percentage, and total area.

## Results

### Comparative Analysis of Patient Tumors (FDTs) and Matched PBT Lines

We assembled and deeply profiled a diverse panel of seven primary GBM and HGG specimens together with their corresponding low-passage PBT spheroid cultures. FDT samples represent the cellular composition of the original surgical specimens, whereas PBT cultures provide patient-specific *in vitro* tumor models enriched for viable tumor cells. The cohort encompassed diverse clinical and molecular characteristics, demonstrating substantial inter-patient genomic heterogeneity (Fig. 1A, Table 1). FDT samples obtained at surgery (ranging from 0.02–12g per sample) and used to generate PBT lines were analyzed by scRNA-seq, multiparameter flow cytometry, and immunohistochemistry, providing a high-resolution reference for tumor composition ^28^.

**Figure 1.**
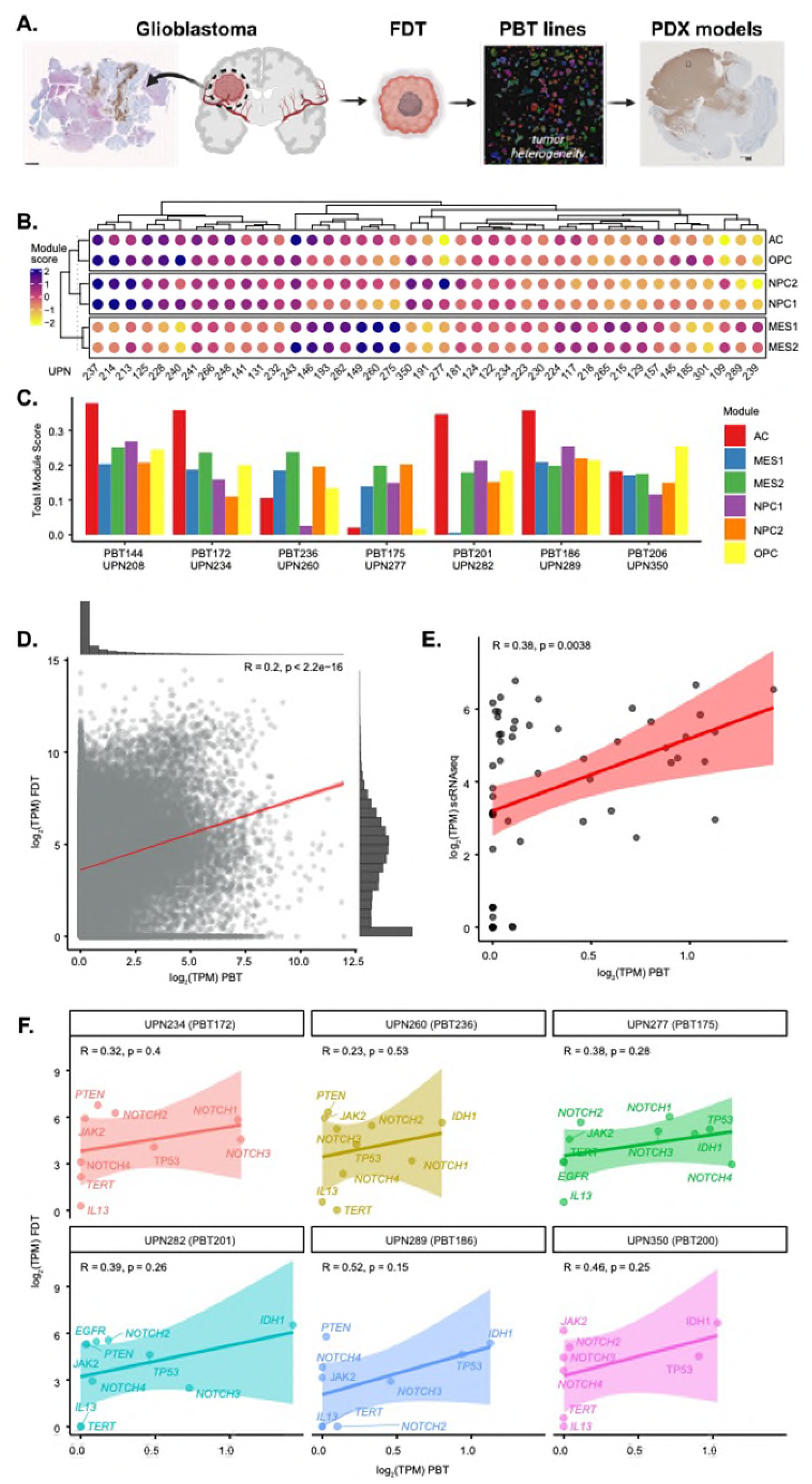
A summary of key genetic alterations and phenotypic features for a panel of PBT lines derived from FDT samples. **(A)** A diagram representing isolation of FDTs, generation of PBT lines, and generation of PDX glioma models. **(B)** GBM meta-module signatures for each FDT sample. Enrichment scores for the degree of over-representation are presented. Scores were calculated using the single-sample gene-set enrichment method (ssGSEA). **(C)** GBM meta-module signatures calculated for each PBT. TotalScore is a combined score that represents the overall expression of a meta-module gene set in a single sample. **(D)** Scatterplot showing gene-level correlation between log₂-transformed TPM (transcripts per million) values from bulk RNA-seq of PBT tissues and their matched FDT samples. Each dot represents a gene. A modest but significant positive correlation was observed (R² = 0.2, *p* < 2.2e–16), suggesting partial transcriptomic fidelity is maintained during initial tumor dissociation. Density plots along the top and right margins depict gene expression distributions in PBT and FDT, respectively. **(E)** Correlation between bulk RNA-seq TPM values from PBT tissues and average expression from corresponding glioma-derived neurospheres assessed by scRNA-seq. A moderate and statistically significant positive correlation was observed (R² = 0.38, *p* = 0.0038), indicating transcriptional continuity between primary tumor tissues and patient-derived *in vitro* neurosphere models. The shaded region indicates the 95% confidence interval for the linear regression. These comparisons validate the relevance of PBT lines derived from FDT samples for functional transcriptomic and therapeutic studies. **(F)** Gene-specific correlation of transcript expression between PBT and FDT samples across individual patients. Scatterplots show the relationship between log₂ TPM values for selected glioma- and immunotherapy-relevant genes in matched PBT and FDT samples across six patient-derived tumor lines (UPNs 234, 260, 277, 282, 289, and 350). Each point represents a gene, including EGFR, PTEN, TP53, IDH1, TERT, JAK2, IL13, and members of the NOTCH family (NOTCH1–4). Regression lines are shown with 95% confidence intervals (shaded). The strength and significance of correlation (Pearson’s R and *p*-value) are indicated for each patient. These analyses highlight the variability in transcriptomic preservation of key oncogenic and immunotherapy targets between PBT and corresponding FDT models. While some samples (e.g., UPN 289, R = 0.52) demonstrate moderate correlation, others show weaker or non-significant associations, reflecting patient-specific transcriptional divergence upon tumor dissociation.

Across all FDT samples analyzed, approximately 68% (37 of 54) successfully formed PBT TSs, enabling downstream modeling of tumor biology. Analysis of the PBT cultures demonstrated that they contained diverse tumor cell populations, indicating that the cultures preserved key aspects of intratumoral heterogeneity from parental tumors. Such heterogeneity is essential for capturing functional differences in therapeutic response during *in vitro* testing. We next performed a comparative molecular analysis of FDTs and their matched PBT lines using scRNA-seq to define driving mutations, tumor-specific transcriptional signatures, and pathway-level alterations (Fig. 1B, C). Each PBT line was characterized alongside its matched FDT counterpart for the expression of actionable oncogenic pathways and known glioma-associated gene signatures (Fig. 1B, C; Fig. S1) ^27^. Bulk RNA-seq meta-module signatures further revealed that PBT samples recapitulated major GBM molecular programs, including proliferative, mesenchymal, and proneural phenotypes (Fig. 1C; Fig. S1, S2).

To quantify molecular fidelity between FDTs and PBTs, we computed global gene expression correlations. Across ten matched pairs, PBT lines demonstrated strong transcriptional similarity to their parent tumors, with an overall correlation coefficient of R = 0.38 (p = 0.0038; Fig. 1D, E). When restricted to curated, glioma-relevant gene sets, correlations increased substantially (Fig. 1E), underscoring the preservation of disease-defining transcriptional programs. Key patient-specific mutations, including *IDH1*, *MGMT*, *TP53*, and *PTEN*, were retained in PBT lines, along with conserved expression of clinically actionable surface and signaling molecules such as IL13Rα2, HER2, EGFR, WNT1, JAK1/2, and NOTCH1-4 (Fig. 1F; Fig. S1, S2). In summary, our findings demonstrate that low-passage PBT lines faithfully preserve the mutational, transcriptional, and pathway-level features of the parent tumors, establishing a robust platform for downstream mechanistic studies and functional therapeutic testing.

### PBT Cell Line Characterization *In Vitro*: Imaging, Doubling Times, Phenotypic Diversity, and Flow Cytometry Profiling

To assess the biological properties of our PBT lines, we performed *in vitro* characterization using imaging-based assessments, proliferation kinetics, and multiparameter flow cytometry. Across the panel of PBTs, we observed substantial variability in growth dynamics, with doubling times ranging from 29 to 160 hours under standardized culture conditions (Table S1). This wide distribution reflects intrinsic biological differences among tumors, including metabolic state, proliferative potential, and degree of stemness, which are features commonly associated with patient-specific glioma heterogeneity. Morphologically, PBT lines demonstrate diverse growth behaviors (Fig. 2A). Cultured under neurosphere-promoting conditions supplemented with EGF and FGF, each line maintained distinct cellular architectures, ranging from loosely aggregated clusters to tightly packed neurospheres or mixed adherent cultures. These morphological differences likely represent underlying transcriptional programs associated with stemness, differentiation gradients, and intrinsic tumor-specific mutational profiles, preserved from the original patient tumors.

**Figure 2.**
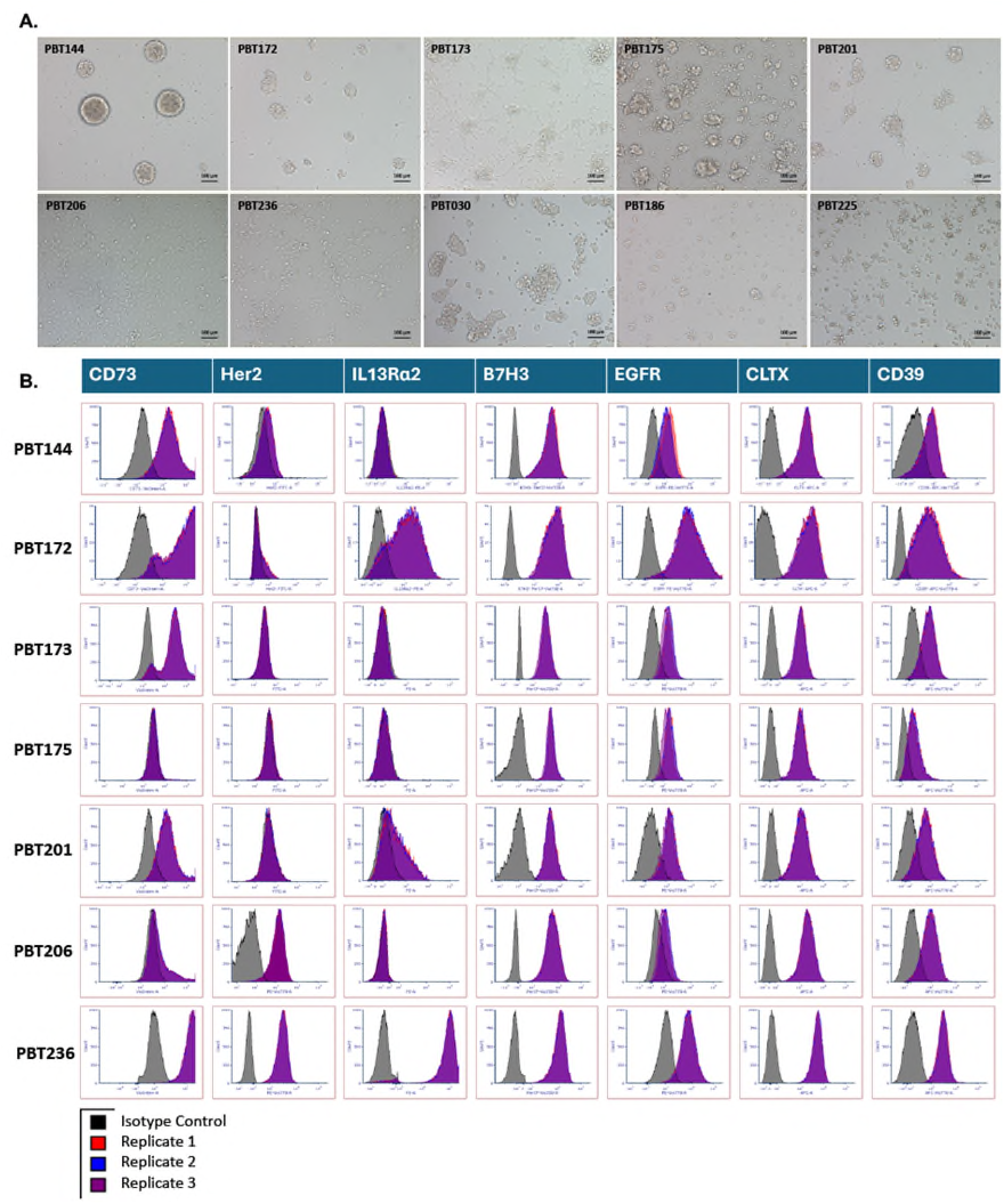
Morphological and immunophenotypic characterization of PBT cell lines. Top panel: Brightfield images of 12 PBT cell lines cultured under neurosphere-forming conditions show varying morphologies, ranging from tight spheroid formation (e.g., PBT144, PBT175) or adherent clusters (e.g., PBT186, PBT225)^43^. Images were taken at 10× magnification; scale bars = 100 µm. The accompanying table summarizes seeding density (cells/cm²), average ± standard deviation, and number of replicates (n) used for growth assessment (table content not shown). Bottom panel: Flow cytometry analysis of surface expression of tumor-associated markers CD73, Her2, IL13Rα2, B7H3, EGFR, CLTX, and CD39 across the 12 PBT lines. Each histogram shows fluorescence intensity of target marker (purple overlay) compared to isotype control (grey). Marker expression varies across tumor lines, highlighting intertumoral heterogeneity in antigen expression relevant for immunotherapy development. These data help inform the selection of appropriate PBT models for preclinical evaluation of targeted therapies such as CAR T cells.

Flow cytometric profiling further revealed that all PBT lines expressed a panel of clinically relevant, targetable surface proteins, though the magnitude of expression varied across samples (Fig. 2B; Fig. S3). Key antigens, including CD73, HER2, IL13Rα2, B7H3, EGFR, CLTX-binding sites, and CD39, were consistently detected, confirming that these models retained surface marker heterogeneity reflective of the patient tumor landscape. Differential expression patterns across the PBT panel suggest that each line may have distinct vulnerabilities to immunotherapies, antibody-drug conjugates, and targeted therapies.

Importantly, the surface biomarker expression observed *in vitro* was highly correlated with that detected in matched PDX tumor xenografts (Fig. 3), underscoring the stability of antigen expression across platforms and validating these PBT lines as biologically relevant preclinical models. This alignment between *in vitro* and *in vivo* systems supports the utility of PBT models for high-throughput screening, mechanistic studies, and preclinical testing of next-generation glioma therapies, including CAR T cells, bispecific antibodies, and other interventions. Together, these findings demonstrate that PBT lines retain key molecular, phenotypic, and proliferative features of the patient tumors from which they were derived, and they highlight the value of these models for dissecting tumor heterogeneity, therapeutic sensitivity, and biomarker-driven treatment stratification.

**Figure 3.**
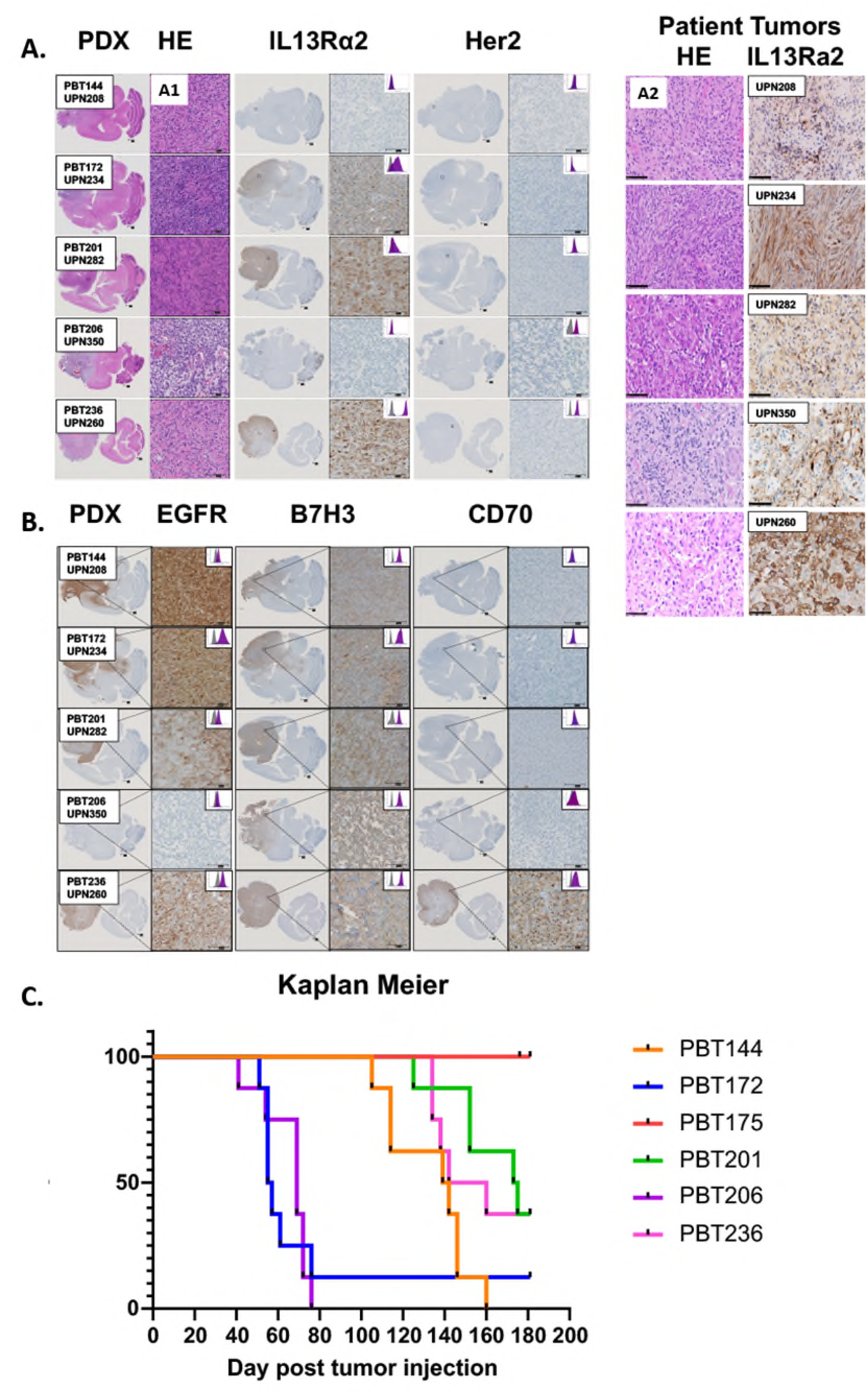
Histopathological and immunohistochemical characterization of PDX brain tumor models and patient tumors. **(A)** Representative sagittal brain sections from NSG mice harboring intracranial xenografts derived from five different PBT lines (PBT154, PBT112, PBT021, PBT206, and PBT236) are shown (A1) and patient tissue (A2). The first column shows hematoxylin and eosin (H&E) staining to assess tumor morphology and infiltration. Adjacent columns display immunohistochemistry (IHC) staining for tumor-associated antigens including IL13Rα2, HER2, EGFR, and B7-H3, highlighting heterogeneous expression across PBT lines. For each marker, low-magnification (left) and high-magnification (right) images are shown to demonstrate regional and cellular expression. Purple box inset diagrams represent the expression of tumor antigens on the corresponding PBT lines as measured by flow cytometry. Staining was performed on formalin-fixed paraffin-embedded (FFPE) brain sections and imaged at both whole-brain and cellular resolution. This panel demonstrates the molecular heterogeneity and preserved expression of immunotherapy-relevant targets across PDX models derived from diverse glioma patients (A1, B), as compared with patient tissue HE and IL13Ra2 staining (A2). **(B)** Kaplan-Meier survival curves of mice bearing intracranial PDX models. NSG mice were stereotactically implanted with tumor cells derived from six distinct PBT lines: PBT144, PBT172, PBT175, PBT201, PBT206, and PBT236. Survival was monitored daily, and animals were euthanized upon the development of neurological symptoms or other humane endpoints. The Kaplan-Meier plot shows variable survival kinetics among the different PDX models, indicating differential tumor aggressiveness and growth rates. Notably, PBT175 and PBT144 exhibited prolonged survival, whereas PBT172 and PBT206 led to rapid disease progression. These models reflect the heterogeneity observed in patient gliomas and are suitable for testing therapeutic responses *in vivo*. Statistical comparison of survival curves was performed using the log-rank (Mantel-Cox) test. All images captured at 10× magnification; scale bars represent 100 microns.

### Orthotopic PDX Models Derived from PBT Lines

To generate orthotopic xenografts, we engrafted PBT lines into the right frontal lobe of NSG mice aged 6–12 weeks and of both sexes. Each mouse received an injection of 0.1 × 10^6^ PBT cells in 2 µL, precisely delivered into the frontal lobe as per established protocols ^32^. Tumor development was closely monitored through daily observations for clinical symptoms such as weight loss, hunching, and changes in behavior or visual acuity, indicative of tumor growth. Upon the manifestation of symptoms consistent with brain tumor progression, mice were humanely euthanized. Their brains were then collected, fixed in 4% paraformaldehyde, and processed by paraffin embedding to prepare for IHC analysis (Fig. 3A). The IHC analysis, conducted using antibodies against key biomarkers (EGFR, IL13Rα2, B7H3, CD70, and Her2), was performed according to the protocols provided by the COH Pathology Core (Fig. 3A, B). The IHC findings were compared with flow cytometry data obtained from the corresponding PBT lines (Fig. 3A), enabling a detailed comparison of biomarker expression between the xenografts and the original PBT lines. Histological examination confirmed that the PBT xenografts faithfully recapitulated the histopathological features of the patient-derived tumors. Additional IHC analysis and HE images were performed for underlying histological features of PDX models and patient samples of glioma (Fig. S4, S5). Kaplan-Meier survival curves were constructed to depict the tumor engraftment, biomarker expression and overall survival of the mice (Fig. 3B), providing insights into tumor growth kinetics in NSG mice, antigen expression and animal survival. Quantitative analysis using QuPath software revealed heterogeneous expression patterns of EGFR and IL13Rα2 across the PDX tumor sections (Table S2). EGFR expression was robust and widespread in several tumor regions, consistent with amplification observed in the corresponding PBT lines. In contrast, IL13Rα2 expression was more spatially restricted and exhibited a patchy distribution, with high expression localized to discrete tumor areas. These findings indicate intertumoral heterogeneity and suggest that EGFR and IL13Rα2 may serve as differential therapeutic targets within the PDX models, as previously described^24^.

### Genomic Characterization of PBT Lines Using OncoScan

To assess chromosomal alterations in FDT samples from individual patients, each designated by a *unique patient number (UPN),* and their corresponding PBT lines, we performed copy number analysis using the OncoScan platform. Genomic profiles of the PBT lines were compared with those of their matched FDT specimens, which had previously undergone targeted whole-genome sequencing (Fig. 4A). The analysis consistently identified hallmark glioma-associated chromosomal alterations, including chromosome 7 amplification and chromosome 10 deletion, across all PBT lines (Fig. 4A). Furthermore, OncoScan revealed variable patterns of copy number alterations and loss of heterozygosity, indicating the presence of both multiple driver mutations and, in some cases, distinct single-gene events within individual lines (Fig. 4B). These findings underscore the genetic complexity and heterogeneity of gliomas and highlight the importance of individualized tumor profiling. This genomic information is critical for identifying tumor-intrinsic features that may influence CAR T cell efficacy and can inform patient-specific immunotherapeutic strategies. By integrating these high-resolution genomic data with functional assays, we aim to establish a robust, scalable platform for precision immunotherapy development.

**Figure 4.**
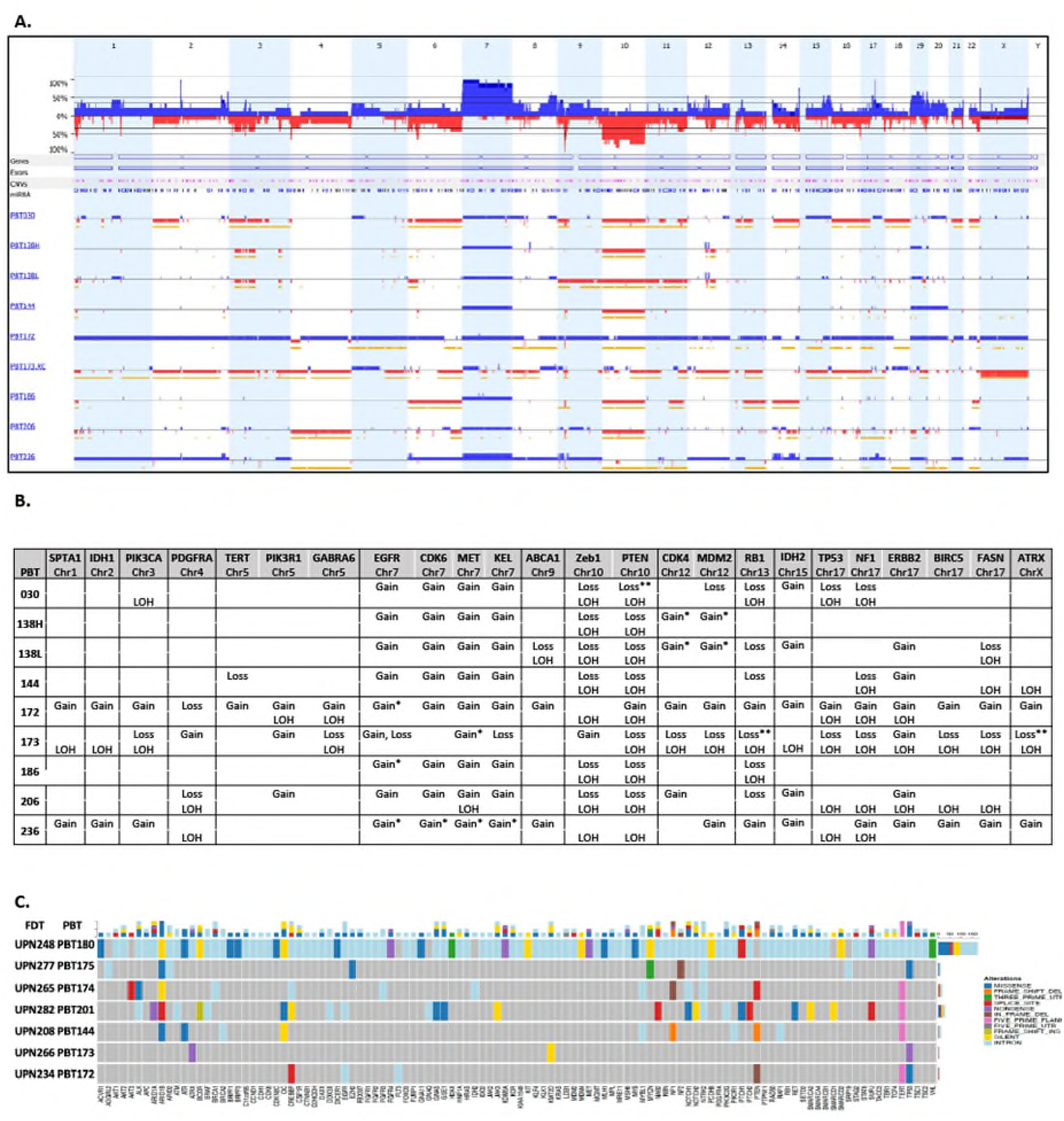
Characterization of Mutational Profiles in PBT Cell Lines and CNV. **(A)** Mutational analysis and chromosomal abnormalities in PBT lines were examined using Oncoscan technology. This analysis identified regions of gain (blue), loss (red), and loss of heterozygosity (yellow) in CNV across the genomes of 9 different PBT lines. The top graph illustrates the percentage of lines that exhibited either gain or loss in each genomic region. **(B)** Panel displaying the specific loss, gain, and loss of heterozygosity (LOH) in 24 key GBM genes. Asterisks denote regions with significant copy number changes: * Indicates high copy gain or loss, while ** indicates homozygous copy loss or gain. **(C) Mutational profile of FDTs (UPN) tumors corresponding to PBTs cell lines**. Summary oncoprint of key genetic alterations and phenotypic features for a panel of FDTs (exome seq). Each row represents an individual GBM patient, and each column corresponds to a gene included in the targeted panel. Colored boxes indicate the presence and type of variant detected in each gene, as defined in the legend on the right. The bar plot at the top summarizes the frequency and type of variant present in that gene across all patient samples, whereas the side bar plot indicates the total number of variants identified per patient. Gray boxes represent genes with no detected alterations.

### Mutational Landscape of GBM Patient Samples was Analyzed by Targeted Sequencing

To characterize gene-level alterations across this GBM cohort, we performed targeted exome sequencing on patient samples and visualized the mutational landscape (Fig. 4C). The analysis revealed marked heterogeneity in mutation number and type detected across patients. Several genes were recurrently altered, including *TP53*, *NF1*, *PTEN*, *TERT*, and *CREBBP*, all of which are well-established drivers of gliomagenesis. Notably, some patients exhibited a broad spectrum of mutations affecting multiple genes (UPN248), while others displayed only a few isolated alterations, reflecting variability in tumor genomic complexity. In addition, we visualized large-scale genomic instability and identified recurrent copy number events in these patient samples using targeted exome sequencing (Fig. S6). The analysis revealed consistent genomic signatures of GBM across multiple samples, including recurrent gains on chromosome 7 and losses on chromosome 10, consistent with our oncoscan results. IDH-mutant tumor (UPN265), clinically classified as an oligodendroglioma, showed 1p/19q-codeletion, consistent with the genetic signature of oligodendroglioma. Additional focal amplifications and deletions were detected on chromosomes 1, 9, and 17, affecting key oncogenes and tumor suppressors such as *CDKN2A/B* (9p), *TP53* (17p), and other GBM-associated genes. Taken together, these findings address the molecular differences of GBM, revealing both recurrently altered driver genes and unique, patient-specific mutations.

### Bulk RNA Sequencing Analysis of PBT Lines

PBT lines underwent bulk RNA sequencing to characterize their transcriptional landscapes. Raw sequencing reads were processed and analyzed using Partek® Flow® (Partek Inc., St. Louis, MO), following the manufacturer’s recommended workflows for alignment, quantification, differential expression, and pathway analysis. Differentially expressed genes were defined using a fold-change cutoff of ±3.5 and a significance threshold of *p* < 0.05, ensuring that only robust transcriptional differences were included in downstream analyses. Principal component analysis (PCA) demonstrated clear separation among PBT lines, indicating substantial inter-line transcriptional heterogeneity and a rightward shift along the primary component axis (Fig. 5A). This variability was further reflected in the heatmap of 23 selected genes, which highlighted distinct expression signatures across PBT models (Fig. 5B).

**Figure 5.**
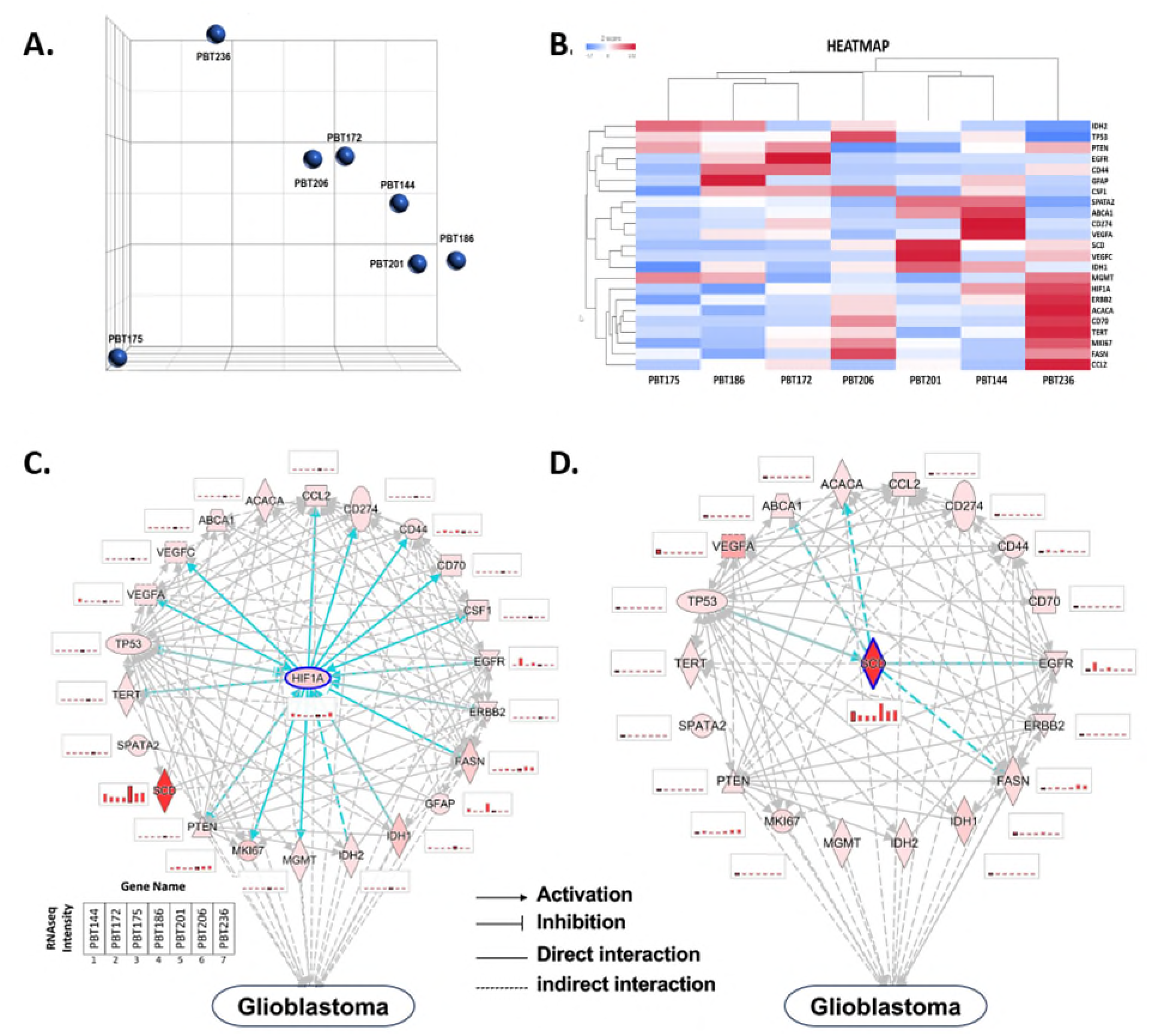
Multi-Omic characterization of PBT lines using PCA, gene expression profiling, and IPA network analysis. **(A)** PCA of seven human glioma PBT tumor cell lines, demonstrating sample clustering based on global bulk RNA-seq gene expression patterns and highlighting transcriptional heterogeneity among PBT models. **(B)** Heatmap of 23 selected genes measured across PBT samples using bulk RNA-seq. Expression patterns reveal inter-tumoral variability in oncogenic, metabolic, and immune-associated genes relevant to GBM biology. **(C)** HIF1A-centered IPA network showing predicted regulatory relationships among selected genes. Nodes represent genes identified in the RNA-seq dataset, and edges denote activation, inhibition, or direct molecular interactions involving the HIF1A signaling axis. **(D)** SCD1-centered IPA network depicting molecular interactions between SCD1 and GBM-associated genes. Bar plots integrated into the network represent relative gene expression (RPKM) values from bulk RNA-seq across the seven glioma PBT lines, illustrating the contribution of SCD1-linked genes to metabolic and oncogenic signaling programs.

Ingenuity Pathway Analysis (IPA) performed on the selected gene set revealed shared regulatory networks across all glioma-derived PBT lines (Fig. 5C, D). Within these networks, HIF1A emerged as a prominent upstream regulator, consistent with its established role in hypoxia-driven glioma biology (Fig. 5C). Leveraging these models, we identified Stearoyl-CoA Desaturase (SCD1) as a novel and consistently overexpressed biomarker across all PBT glioma lines and matched FDT patient tumors. IPA revealed SCD1 as an upstream regulator of key oncogenic and metabolic pathways, including EGFR, TP53, and CYCS, thus supporting its role in glioma progression and in shaping an immunosuppressive tumor microenvironment (TME) (Fig. 5D). Collectively, these analyses indicate that PBT lines retain distinct and biologically relevant transcriptional programs and identify SCD1 as a novel, consistently elevated biomarker with mechanistic links to metabolic and oncogenic networks in glioma.

### SCD1 Expression Landscape Across FDT Tumors, Gene Networks, Clinical Outcomes, and CAR T Therapy Response

Single-cell transcriptional profiling of all UPN samples derived from patient samples (FDTs) of patients participating in CAR T clinical trial (COH IRB13384) revealed distinct malignant and non-malignant cell populations, including multiple tumor clusters, oligodendrocyte-lineage cells, myeloid subsets, lymphocytes, and fibroblasts (Fig. 6A). Mapping SCD1 expression onto this cellular landscape demonstrated that SCD1 is selectively enriched within malignant tumor clusters and largely absent from non-tumor compartments, indicating a tumor-specific metabolic signature in scRNA-seq analysis of FDTs (Fig. 6B). Parallel analysis of PBT lines showed consistent upregulation of SCD1 across low passage tumor lines, confirming that elevated SCD1 expression is maintained *in vitro*, not only in the original tumor tissue (Fig. 6C).

**Figure 6.**
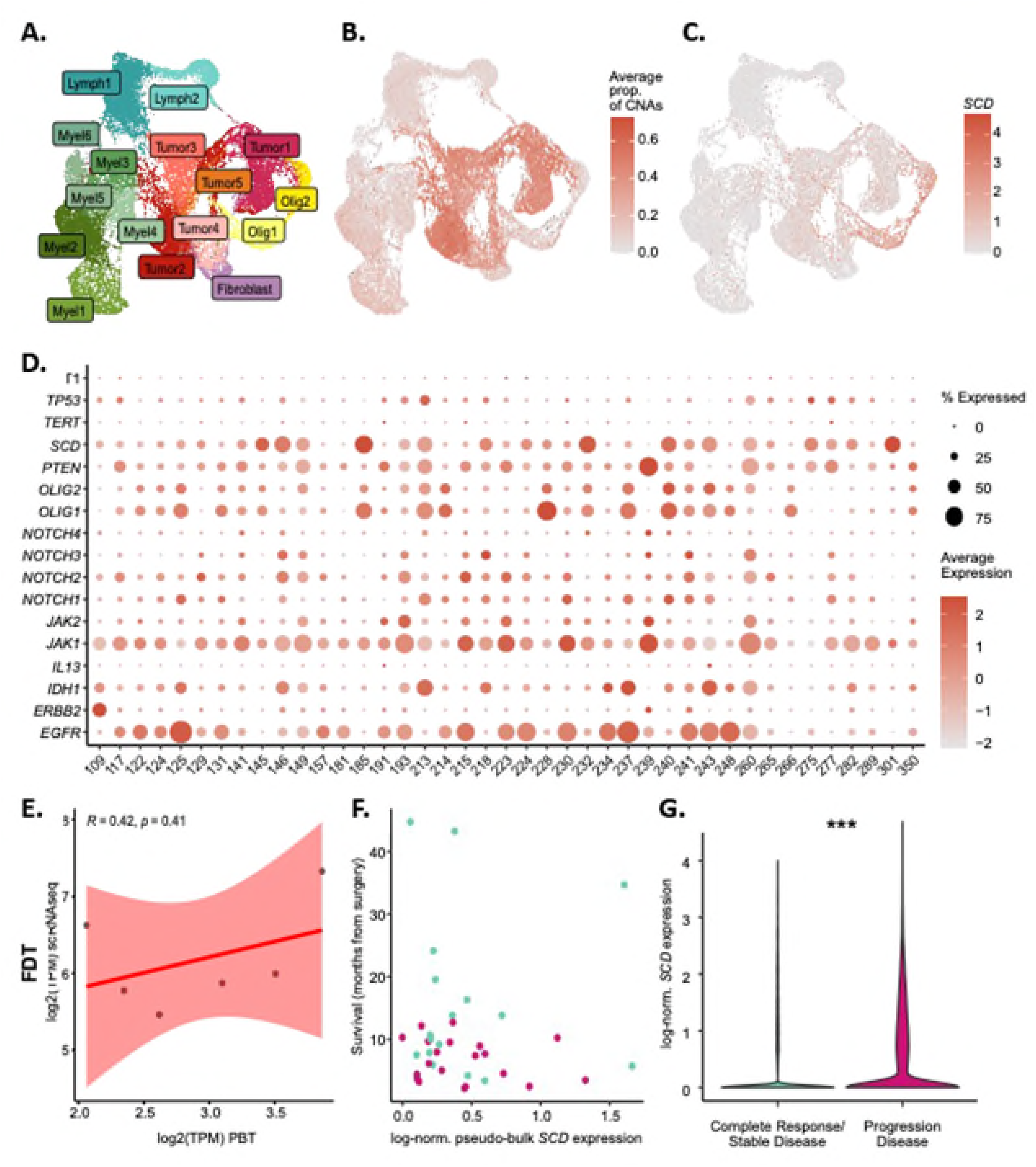
SCD1 expression landscape across patient tumors, PBT lines, gene networks, TME composition, survival outcomes, and CAR T therapy response. **(A)** UMAP projection of all UPN samples, annotated by major cell populations, including tumor clusters, oligodendrocyte-lineage cells, myeloid subpopulations, lymphocytes, and fibroblasts, illustrating transcriptional architecture across patient tumors. **(B)** SCD1 expression across all UPN cells, demonstrating preferential enrichment of SCD1 within malignant tumor clusters relative to non-malignant populations. **(C)** SCD1 expression across PBT lines, showing conserved and enhanced upregulation of SCD1 in patient-derived tumor cell lines compared with matched UPN samples. **(D)** Dot plot of SCD1-correlated genes across UPNs and PBTs, highlighting co-expression networks associated with metabolic regulation, apoptosis (including CYCS), lipid biosynthesis, oncogenic signaling (EGFR, NOTCH, TP53 pathways), and immune-modulatory programs. **(E)** Distribution of SCD1 expression across tumor and stromal cell types, demonstrating relative contributions of fibroblasts, myeloid cells, and other TME components to the overall SCD1 transcriptomic signature. **(F)** Correlation of SCD1 expression with clinical outcomes, showing that high SCD1 expression is associated with poorer survival trajectories, including increased frequency of progressive disease (PD) compared with stable disease (SD) across the patient cohort. **(G)** SCD1 expression as a biomarker of CAR T cell therapy response, where SCD1-high tumors exhibit distinct response patterns and reduced sensitivity to CAR T–mediated tumor control, suggesting a mechanistic link between SCD1-driven metabolic reprogramming, immune suppression, and therapeutic resistance.

To define the broader transcriptional programs associated with SCD1 activity, we examined its correlation with a curated gene panel relevant to glioma metabolism, oncogenic signaling, hypoxia, immune modulation, and therapeutic resistance. Dot-plot visualization revealed strong positive associations between SCD1 and genes involved in lipid metabolism (FASN, ACACA), apoptosis regulation (CYCS), oncogenic drivers (EGFR, ERBB2, TERT), and immune-suppressive mediators (CD44, CD274/PD-L1, CCL2) (Fig. 6D). These patterns suggest that SCD1 integrates metabolic reprogramming with oncogenic pathways across patient tumors. Our study is in line with recently reported identification of SCD15 (stearoyl-CoA desaturase-5) as a regulator of GBM stem cell (GSC) survival and genomic integrity^38^. Although SCD1 has been widely recognized as an important lipid metabolic enzyme in GBM, the findings demonstrate that SCD5 performs unique functions that cannot be compensated by SCD1, highlighting its specific importance in revealing a direct connection between fatty acid desaturation, lipid homeostasis, and DNA repair pathways in brain tumors. Decomposition of TME composition demonstrated that SCD1-high tumors exhibit distinct cellular proportions, including increased fibroblast and myeloid cell representation (Fig. 6E), consistent with a more immunosuppressive and stromal-enriched TME state. Clinically, elevated SCD1 expression was associated with poorer outcomes, including a higher likelihood of progressive disease (PD) relative to stable disease (SD) among treated patients in CAR T clinical trial (NCT02208362) (Fig. 6F). This trend suggests that SCD1 may serve as a prognostic biomarker reflecting more aggressive tumor behavior, linking tumor metabolism and DNA repair pathways^38^. Finally, when SCD1 expression was examined in the context of CAR T cell therapy responses, tumors with high SCD1 levels demonstrated reduced therapeutic benefit, with diminished or absent response signatures compared to SCD1-low tumors (Fig. 6G). These findings highlight SCD1 as a potential biomarker of CAR T resistance and implicate metabolic remodeling in shaping the immunosuppressive environment that constrains T cell efficacy.

Together, these data position SCD1 as a central metabolic–immune regulator that defines tumor identity, correlates with oncogenic and immune-modulatory gene programs, influences TME composition, predicts clinical outcomes, and may inform response to CAR T cell therapy in glioma.

## Discussion

Previous studies have established large collections of patient-derived GBM cultures, organoids, and xenografts that retain key molecular and phenotypic characteristics of the original tumors and serve as valuable platforms for translational research and therapeutic testing^39^. Notably, cohorts such as the Human Glioma Cell Culture (HGCC) collection and other pharmacogenomically annotated patient-derived glioma panels have provided important insights into intertumoral heterogeneity and treatment response^40, 41^. A particular strength of the present PBT collection is its well-characterized panel of patient-derived models that capture clinically relevant tumor diversity while providing a reproducible experimental resource for functional studies. In this regard, the collection complements existing glioma biobanks by enabling systematic investigations of tumor biology and therapeutic vulnerabilities in models that closely reflect patient tumors^42^.

Our comprehensive genomic profiling of PBT lines using OncoScan technology revealed recurrent chromosomal alterations frequently observed in HGGs, including chromosome 7 gain and chromosome 10 loss. These aberrations are well-characterized hallmarks of GBM multiforme in large consortia such as The Cancer Genome Atlas (TCGA), which are associated with alterations of key oncogenes and tumor suppressors, such as **EGFR**, **MET**, **PTEN**, and **CDKN2A**. The consistent detection of these genomic events across our low-passage PBT lines affirms their ability to recapitulate the core genetic features of the primary tumors from which they were derived.

In addition to these canonical alterations, OncoScan analysis uncovered diverse patterns of copy number variation and widespread regions of loss of heterozygosity (LOH), highlighting the intertumoral heterogeneity that defines glioma biology. These unique and shared genomic signatures across the PBT cohort suggest that, while certain oncogenic drivers are conserved, many tumors harbor private alterations that may modulate therapeutic susceptibility and resistance. This observation aligns with recent single-cell and spatial transcriptomic studies that have revealed GBM to be made of subclonal populations with distinct proliferative, stem-like, and immunosuppressive phenotypes within the same tumor, identifying mesenchymal, proneural, and OPC signatures within FDTs and PBTs. Thus, our panel of PBT lines offers a clinically relevant platform that captures both convergent evolution (shared drivers) and divergence (private subclones), making it ideally suited for precision modeling of therapeutic response.

Beyond their genomic integrity, these PBT lines are annotated with detailed clinical metadata, including patient age, sex, tumor subtype, and treatment history, allowing for multidimensional analyses that integrate clinical and molecular features. When paired with bulk and single-cell transcriptomic profiling, flow cytometry, and immunohistochemistry, this dataset becomes a powerful tool for dissecting tumor-intrinsic factors that influence therapy response. For example, expression levels and mutational status of targets such as IL13Rα2, HER2, EGFR, JAK1/2, and NOTCH1–4 are not only preserved in our models but can be analyzed in the context of patient outcome and treatment exposure. This integrative approach facilitates the identification of candidate biomarkers and resistance mechanisms relevant to ongoing and emerging clinical trials, particularly in the immunotherapy space.

In our study, SCD1 emerged as a uniformly upregulated biomarker across all PBT glioma lines, highlighting its potential role in glioma pathophysiology. SCD1 is a central enzyme in lipid metabolism, responsible for generating monounsaturated fatty acids that regulate membrane fluidity, organelle function, and oncogenic signaling. Its consistent elevation in glioma cells likely reflects a metabolic adaptation that promotes proliferation, survival under hypoxic stress, and resistance to metabolic constraint. IPA network analysis positioned SCD1 upstream of key oncogenic regulators, including EGFR and TP53, suggesting that its activity may influence broader tumor-promoting signaling networks. The identification of SCD1 as a central molecular node not only underscores its biological relevance but also raises the possibility that metabolic targeting of SCD1 could disrupt multiple convergent pathways critical for glioma progression. Given its robust expression across models, SCD1 may represent both a biologically meaningful classifier and an actionable metabolic target in glioma^41^.

Importantly, our findings demonstrate that low-passage PBT lines and matched PDXs preserve heterogeneous, non-clonal cellular architectures, even under *in vitro* and *in vivo* selection pressures. This maintenance of intratumoral heterogeneity is essential for modeling therapeutic resistance, as clonal and subclonal adaptation drive treatment failure in glioma. Prior studies have shown that CAR T cells, immune checkpoint inhibitors, and kinase inhibitors often select for resistant populations through antigen downregulation, epitope alteration, or activation of compensatory signaling circuits. Our models provide a tractable system to study these evolutionary dynamics in real time, enabling functional interrogation of resistance mechanisms and evaluation of rational combination therapies designed to limit clonal escape.

The implications for immunotherapy development are substantial. Tumor-intrinsic genomic and transcriptomic features, such as antigen heterogeneity, PTEN loss, impaired interferon signaling, and altered MHC expression, shape response to CAR T cells and other immune-based modalities. By integrating multi-omic profiling with *in vitro* CAR T co-culture assays and *in vivo* PDX testing, we can stratify PBT lines based on predicted immunotherapy responsiveness. This platform allows for a detailed understanding of immune resistance, including spatial immune exclusion, cytokine-driven immunosuppression, and transcriptional programs associated with T cell exhaustion. The establishment of a well-characterized library of PBT lines and corresponding PDX models provides a clinically relevant platform for evaluating CAR T-cell immunotherapy and identifying patient-specific therapeutic sensitivities. Although generation of stable PBT lines typically requires 2–4 months, development of rapid 3D spheroid-based assays from patient tumors could enable parallel, near-real-time testing of responses to CAR T cells and other therapeutic agents. Integration of these functional drug-response data with molecular tumor profiling may ultimately support individualized treatment selection and provide a translational framework for incorporating patient-specific therapeutic sensitivity testing into clinical decision-making.

In summary, our study establishes patient-derived brain tumor models as a translational bridge between molecular characterization and functional therapeutic testing. The ability of low-passage PBT cultures and matched PDX models to preserve clinically relevant tumor features supports their use for interrogating treatment response in a patient-specific context. When coupled with rapid 3D spheroid assays and molecular profiling, this platform could facilitate prospective assessment of sensitivity to CAR T-cell therapies and other emerging treatments, while helping identify mechanisms associated with therapeutic response or resistance. Ultimately, such integrated functional modeling may contribute to more informed treatment selection and the development of individualized therapeutic strategies for patients with brain tumors.

## Acknowledgements

This work was supported by The Harriet H. Samuelsson Foundation (Margarita Gutova and Christine Brown) and Merkin Family Foundation (Christine Brown). The authors thank Dr. Erin S. Keebaugh for her expert scientific writing and editorial assistance in the preparation of this manuscript. Her contributions to improving the clarity, organization, and presentation of the manuscript are gratefully acknowledged.

