## Supplemental Table and Figures for "Patient-Derived Glioma Models Preserve Tumor Heterogeneity and Identify Stearoyl-CoA Desaturase1 (SCD1) as a Candidate Biomarker for Precision Immunotherapy"

A.

B.

C.

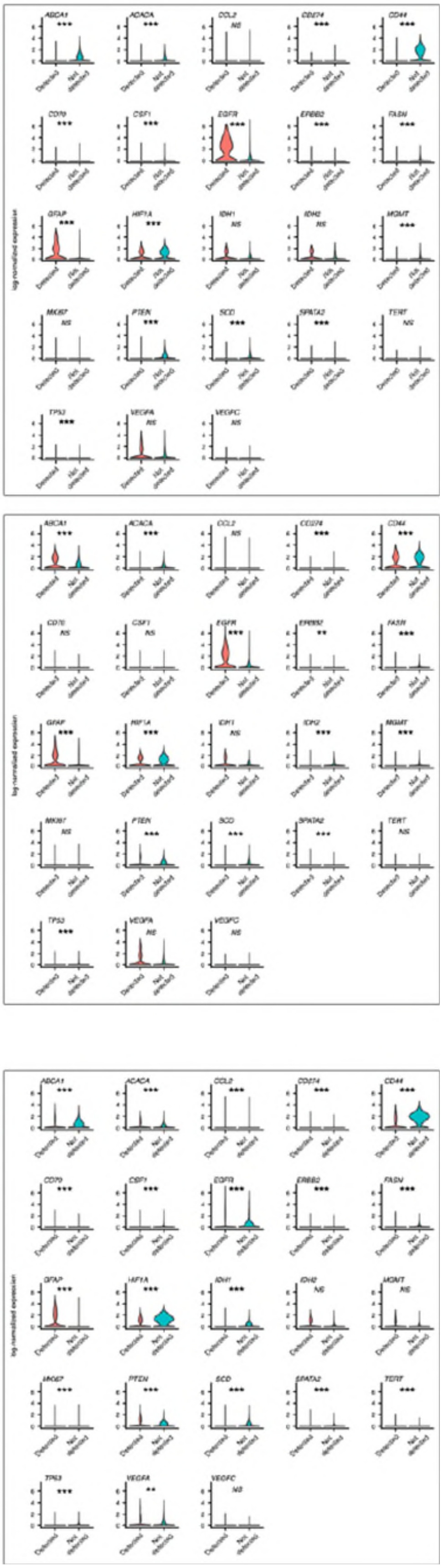

**Figure S1.** Expression analysis of selected glioma-relevant genes classified by EGFR, PTEN, and TP53 mutation status within freshly dispersed tumors (FDTs).

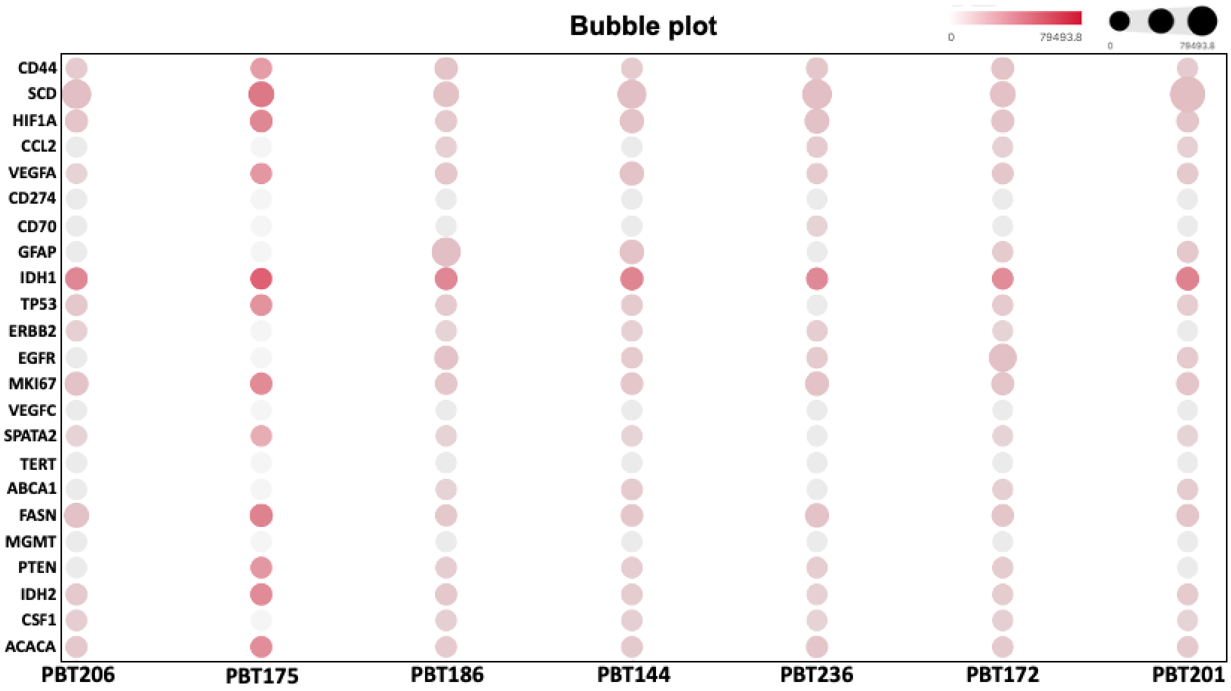

**Figure S2.** Expression analysis of selected glioma-relevant genes in PBT cell lines.

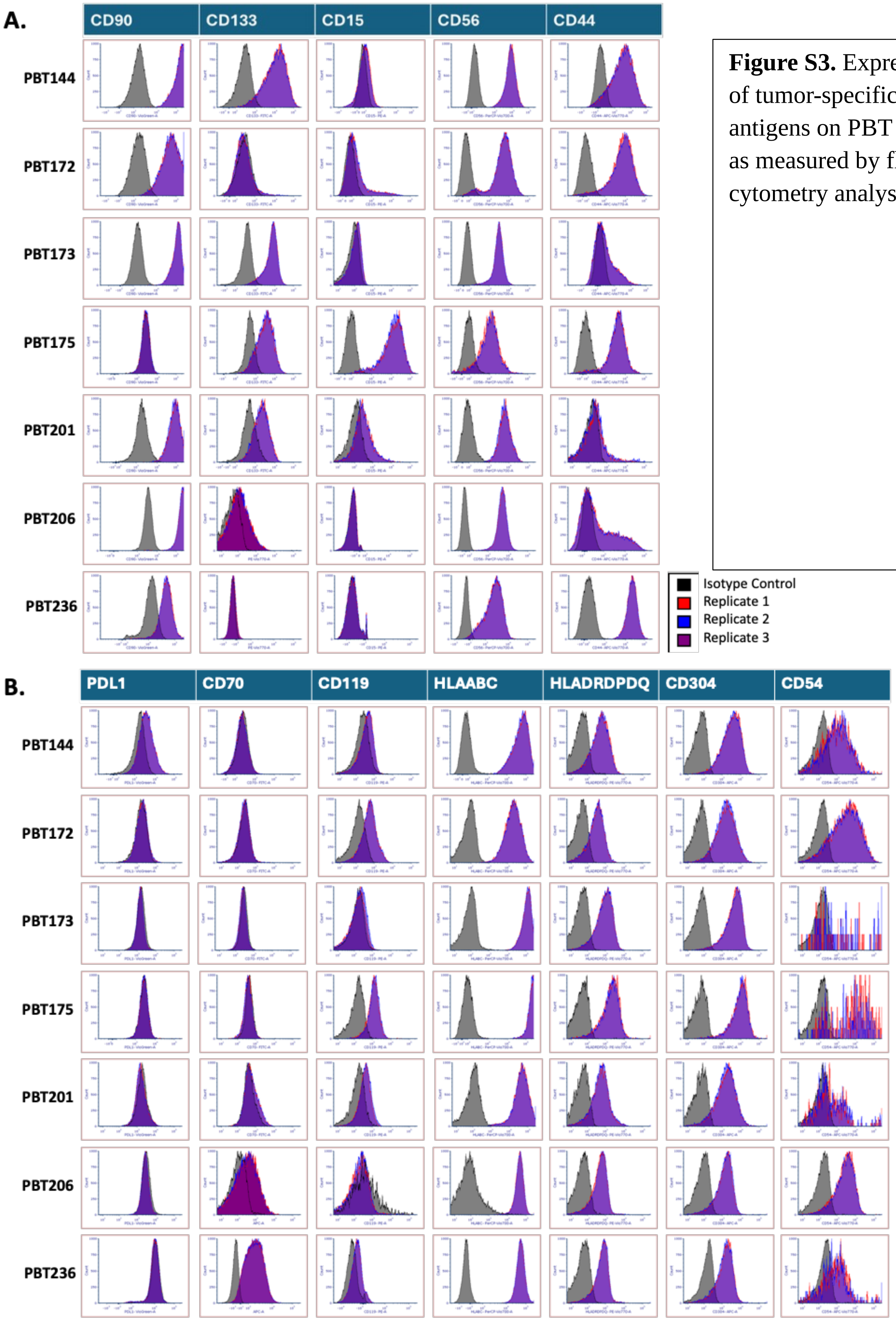

**Figure S3.** Expression of tumor-specific antigens on PBT lines, as measured by flow cytometry analysis.

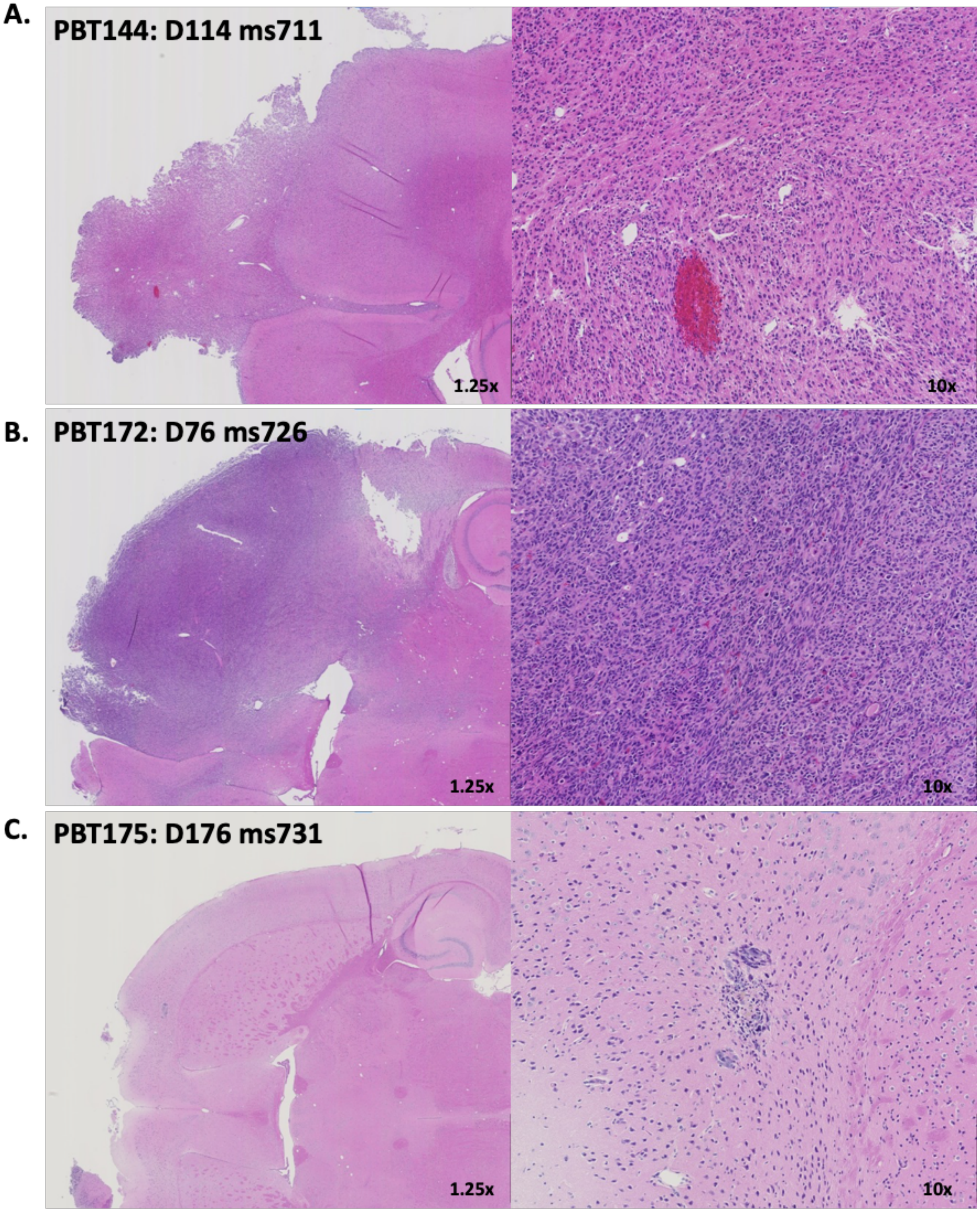

**Figure S4.** Representative hematoxylin and eosin (H&E) stained sections of PDX mouse xenografts.

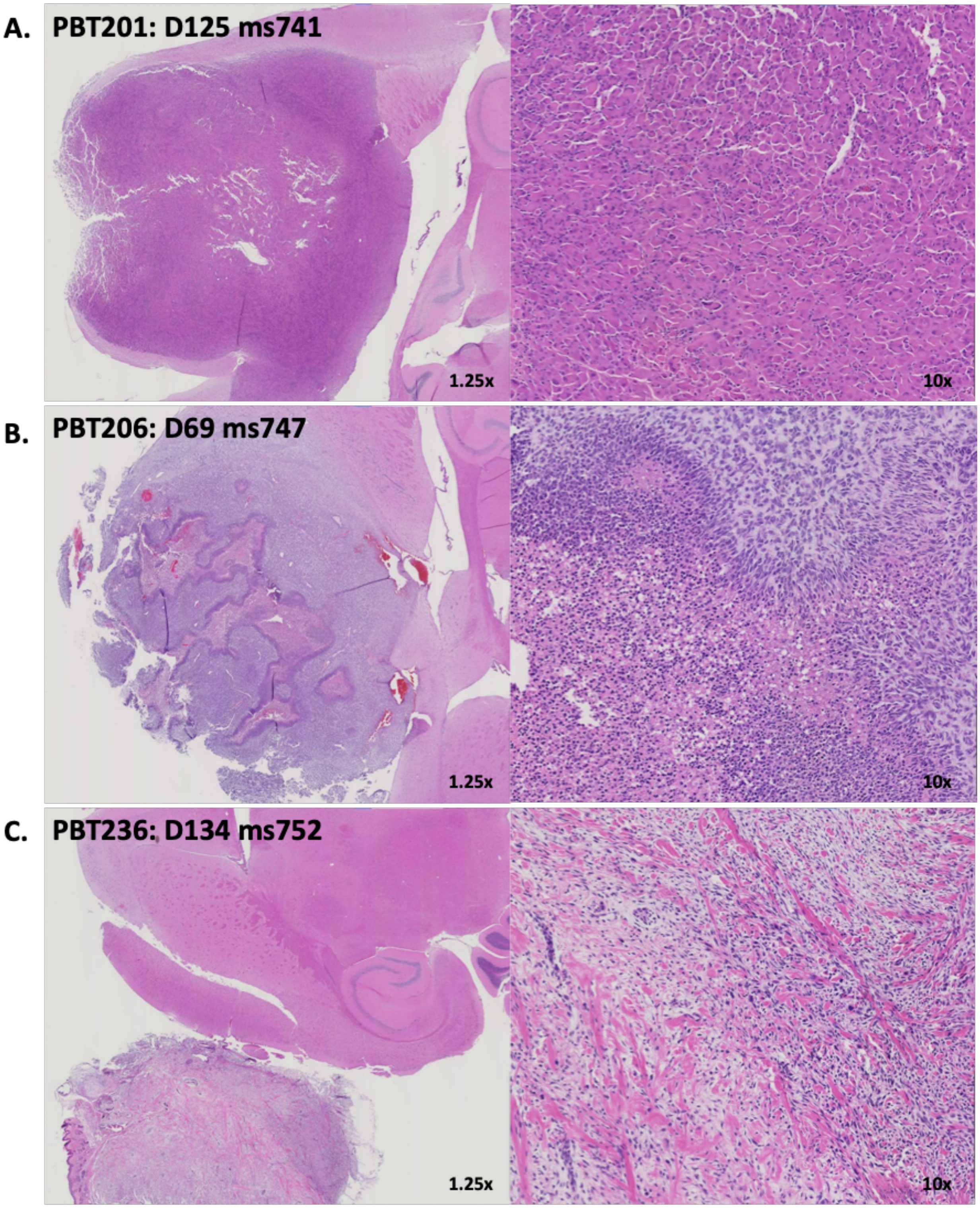

**Figure S5.** Representative hematoxylin and eosin (H&E) stained sections of PDX mouse xenografts.

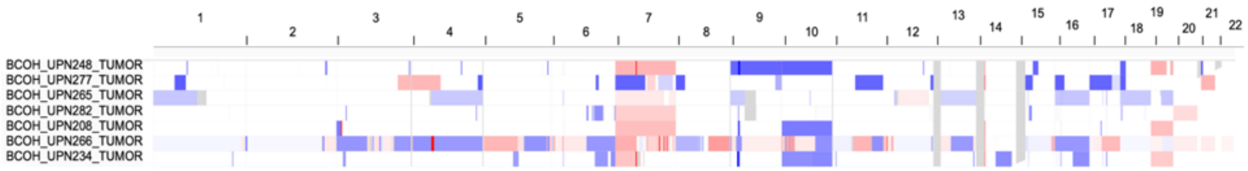

**Figure S6. Copy number alteration profiles of FDT tumors.** Genome-wide copy number alteration landscape of FDT samples (UPNs) visualized in Integrative Genomics Viewer (IGV). Each track corresponds to a single tumor sample, with regions of copy number gain displayed in **red** and regions of copy number loss displayed in **blue**.

**Table S1. PBT line cell culture and doubling time.**

| PBT | Seeding Density (Cells/cm2) | Average ± Standard Deviation | n |
|---|---|---|---|
| 144 | 5.33E+04 | 67.2 ± 37.2 | 3 |
| 175 | 4.00E+04 | 59.1 | 1 |
| 186 | 6.67E+04 | 103.4 ± 67.4 | 2 |
| 201 | 4.53E+04 | 90.1 | 1 |
| 172 | 5.33E+04 | 74.4± 24.87 | 5 |
| 173 | 5.33E+04 | 83.5 ± 11.1 | 2 |
| 206 | 2.67E+04 | 32.9 ± 3.1 | 7 |
| 216 | 2.67E+04 | 29.3 ± 3.2 | 7 |
| 030 | 4.00E+04 | 48.7 ± 8.6 | 3 |
| 138H | 4.00E+04 | 57.1 ± 11.6 | 2 |
| 225 | 2.67E+04 | 86.9± 4.1 | 2 |

### Table S2. Expression of tumor-specific antigens on PDX tumor xenograft models.

| | PBT144 | | | PBT172 | | | PBT173 | | | PBT175 | | | PBT201 | | | PBT206 | | | PBT236 | |
|---|---|---|---|---|---|---|---|---|---|---|---|---|---|---|---|---|---|---|---|---|
| | AVG | SD | | AVG | SD | | AVG | SD | | AVG | SD | | AVG | SD | | AVG | SD | | AVG | SD |
| CD73 | 80.51 | 0.75 | | 86.57 | 0.05 | | 84.04 | 0.44 | | 4.47 | 0.29 | | 65.03 | 0.38 | | 22.30 | 0.25 | | 93.41 | 0.08 |
| Her2 | 7.19 | 0.24 | | 8.06 | 0.59 | | 0.71 | 0.07 | | 2.35 | 0.40 | | 1.02 | 0.00 | | 0.19 | 0.26 | | 1.64 | 0.13 |
| IL13Ra2 | 0.71 | 0.07 | | 73.30 | 0.98 | | 0.35 | 0.06 | | 1.57 | 0.14 | | 80.63 | 0.53 | | 1.12 | 0.55 | | 93.86 | 0.11 |
| B7H3 | 95.75 | 0.04 | | 99.75 | 0.07 | | 99.91 | 0.01 | | 95.54 | 0.13 | | 99.97 | 0.01 | | 99.04 | 0.02 | | 99.90 | 0.03 |
| EGFR | 28.46 | 9.56 | | 94.60 | 0.95 | | 29.37 | 10.59 | | 72.93 | 12.12 | | 55.59 | 7.13 | | 6.97 | 2.12 | | 89.72 | 1.86 |
| CLTX | 96.47 | 0.28 | | 98.93 | 0.32 | | 99.16 | 0.15 | | 99.02 | 0.11 | | 98.53 | 0.36 | | 59.72 | 0.51 | | 99.94 | 0.05 |
| CD39 | 44.03 | 1.59 | | 97.13 | 1.57 | | 48.36 | 2.24 | | 29.47 | 2.01 | | 39.04 | 0.49 | | 63.90 | 1.42 | | 96.07 | 0.14 |
| CD90 | 95.52 | 0.04 | | 95.41 | 0.12 | | 99.50 | 0.01 | | 0.75 | 0.19 | | 95.61 | 0.10 | | 98.92 | 0.20 | | 53.09 | 0.06 |
| CD133 | 87.01 | 0.47 | | 2.89 | 0.23 | | 91.52 | 0.19 | | 66.30 | 0.60 | | 80.04 | 1.79 | | 0.31 | 0.04 | | 1.11 | 0.16 |
| CD15 | 4.17 | 0.88 | | 16.42 | 0.26 | | 0.82 | 0.04 | | 97.84 | 0.06 | | 31.95 | 1.15 | | 0.31 | 0.04 | | 0.75 | 0.02 |
| CD56 | 99.61 | 0.04 | | 94.17 | 0.07 | | 99.01 | 0.00 | | 94.74 | 0.50 | | 99.91 | 0.04 | | 99.71 | 0.22 | | 94.95 | 0.07 |
| CD44 | 83.67 | 0.59 | | 94.55 | 0.01 | | 21.47 | 0.45 | | 93.56 | 0.09 | | 7.53 | 0.30 | | 37.34 | 0.53 | | 99.74 | 0.06 |
| PDL1 | 15.82 | 0.24 | | 1.15 | 0.01 | | 0.35 | 0.05 | | 1.05 | 0.43 | | 1.73 | 0.31 | | 0.44 | 0.52 | | 0.47 | 0.06 |
| CD70 | 0.99 | 0.05 | | 1.09 | 0.10 | | 0.33 | 0.03 | | 0.79 | 0.15 | | 0.29 | 0.15 | | 0.10 | 0.02 | | 1.49 | 0.03 |
| CD119 | 4.47 | 0.21 | | 17.67 | 0.38 | | 1.08 | 0.45 | | 72.34 | 3.57 | | 12.73 | 2.41 | | 1.01 | 0.04 | | [illegible] | 0.22 |
| HLAABC | 95.02 | 0.03 | | 95.88 | 0.02 | | 99.94 | 0.01 | | 96.30 | 0.00 | | 95.93 | 0.31 | | 59.76 | 0.55 | | 99.94 | 0.04 |
| HLADRDP DQ | 55.12 | 0.47 | | 50.11 | 0.01 | | 66.27 | 0.03 | | 75.34 | 0.14 | | 60.52 | 0.74 | | 65.00 | 1.51 | | 73.76 | 1.06 |
| CD304 | 91.41 | 0.07 | | 93.61 | 0.49 | | 79.48 | 1.29 | | 90.48 | 0.62 | | 66.79 | 0.47 | | 64.01 | 0.80 | | 57.31 | 1.20 |
| CD34 | 6.57 | 0.00 | | 37.30 | 0.28 | | 0.17 | 0.04 | | 1.75 | 0.31 | | 1.03 | 0.34 | | 41.59 | 0.30 | | 1.51 | 0.00 |
| VEGFA | 93.31 | | | 80.65 | | | 98.59 | | | 58.7 | | | 75.07 | | | 98.47 | | | 93.15 | |
